# Multi-resolution QTL mapping in four interconnected rice RIL populations reveals recurrent, background-specific, and environment-dependent loci

**DOI:** 10.64898/2026.11.23.768548

**Authors:** Claudio Brondani, Daniany Adorno Rodrigues Silva, Ana Letycia Basso Garcia, Jéssica Fernanda Ferreira dos Santos Prado, Antônio Carlos Centeno Cordeiro, Rosana Pereira Vianello

## Abstract

We aimed to determine how marker density influences the detection and localization of quantitative trait loci (QTLs) in rice and to identify parental alleles with consistent effects across genetic backgrounds and environments. We evaluated grain yield, plant height, and flowering time in four interconnected recombinant inbred line populations at two Brazilian sites, Goianira and Boa Vista. Each parent contributed to two crosses, allowing us to compare the direction of its allelic effects in different genetic backgrounds. We compared a common backbone of 2,060 DArTseq SNPs with denser, cross-specific panels using marker and bin regression, interval mapping based on genotype probabilities inferred by hidden Markov models, and genome-wide permutation thresholds. The backbone recovered the major QTL regions. Peak positions differed by less than 100 kb in 14 of 19 comparisons within crosses and environments, including five exact matches. Across the four maps, densification increased marker counts 4.66-fold and recombination bin counts 2.44-fold. Recombination-breakpoint interval width decreased by 31.1% in Maninjau × Epagri 108 and 90.3% in Irat 122 × Araguaia. Comparisons using the same lines resolved an apparent peak shift at qPH12 and placed qYLD2 below the genome-wide threshold in both panels. Four QTL regions recurred across crosses. Epagri 108 contributed all of them for reduced height at qPH1, whose interval contains *SD1*, and delayed flowering at qDTH8. Araguaia contributed an allele for earlier flowering at qDTH3 in both its crosses. The qDTH6 region recurred in Goianira, and the opposite flowering effects observed across environments in Irat 122 × Araguaia mapped to different chromosome 6 peaks. Grain-yield QTLs showed less recurrence. The strongest associations, qYLD1 and qYLD6, occurred in separate *indica* × *japonica* crosses. In Goianira, qYLD6 explained 32.5% of phenotypic variance in the full-panel analysis and achieved genome-wide significance only at that site. Parental genome sequences helped prioritize candidate genes. Two complementary recombinants delimited a 247.0-kb region containing *Ef-cd* at qDTH3, while *Hd1* and *OsCLF* remained genetically unresolved within a 51.8-kb marker-defined interval at qDTH6. These findings identify Epagri 108 × Araguaia as a candidate cross for combining reduced height with earlier flowering. For grain yield, the favorable Maninjau allele at qYLD1 and Epagri 108 allele at qYLD6 warrant testing individually and together in recipient backgrounds under target growing conditions. Because the backbone was derived retrospectively, its economic benefit requires prospective evaluation. For routine marker-assisted selection, breeders could genotype founders and initial mapping families at high density, use a common panel of markers representing recombination bins to track validated haplotypes across subsequent families, and add cross-specific flanking markers to select progeny that retain a favorable allele within a smaller donor segment.

## INTRODUCTION

QTL mapping in biparental populations remains one of the most direct approaches for dissecting the genetic architecture of complex traits in plants. Recombinant inbred lines (RILs) are particularly useful because they fix combinations of parental alleles, allow for replicated evaluations across different environments, and provide a renewable genetic resource for subsequent validation and fine mapping (Bazakos et al., 2017). However, the interpretation of a QTL depends on the genetic background in which it segregates, the quality of the phenotypic data, the actual amount of observed recombination, and the density and quality of the available markers (Bazakos et al., 2017). Using multiple related populations broadens this perspective. Populations that share common parents allow researchers to investigate whether a specific chromosomal region shows consistent effect directions and comparable effect magnitudes across different backgrounds, or whether its behavior depends on specific allelic combinations and the environment (Palanog et al., 2023; Li et al., 2024). In rice, multi-population designs have improved mapping precision and enabled the estimation of specific parental effects (Fragoso et al., 2017). Nevertheless, direct comparison requires a harmonized genotypic framework; marker sets specific to individual crosses may differ in terms of density, informativeness, missing data, and allele frequencies.

Increasing marker density does not translate linearly into an increase in information. Co-segregating SNPs belong to the same recombination unit; treating them as independent events can lead to the expansion of genetic maps or the amplification of errors (N’Diaye et al., 2017). In maps that are already near saturation, very dense marker sets may not necessarily increase detection power compared to maps of intermediate density, although they do improve the physical localization of crossovers and facilitate fine mapping (Li et al., 2015). Therefore, the most appropriate unit for comparing marker density is the “recombination bin,” rather than the raw number of SNPs.

The experimental set described here comprises four RIL (recombinant inbred line) populations organized into a partial circular diallel mating design. Maninjau and Epagri 108 belong to the *indica* group, while Irat 122 and Araguaia belong to the *japonica* group. Each parent is involved in two crosses, creating a network that encompasses *indica* × *japonica*, *indica* × *indica*, and *japonica* × *japonica* combinations. The populations were evaluated for yield, plant height, and flowering time at two locations—Goianira and Boa Vista—enabling the simultaneous investigation of allelic recurrence, background specificity, and QTL-by-environment interactions. An initial analysis was conducted using 2,060 consensus SNPs common to all four crosses. Subsequently, each population was re-analyzed using the full set of available DArTseq SNPs, resulting in four “full-panel” datasets. This structure provides a unique opportunity to assess sensitivity: the common panel ensures cross-population comparability, while the full panel tests the robustness of the genetic architecture, refines marker positions, reveals previously undetected crossover events, and may also uncover subtle or “fragile” signals.

The objective of this study was to integrate data from four RIL populations evaluated at two locations, conducting a multi-resolution analysis of QTLs related to yield, plant height, and flowering time. Specifically, we aimed to: (i) quantify gains in terms of markers, genomic bins, and breakpoint resolution; (ii) evaluate the recovery of QTLs originally detected using the 2,060-SNP backbone; (iii) distinguish between robust, reclassified, background-specific, and environment-dependent QTLs; (iv) identify recurrences associated with shared parental lines; and (v) establish priorities for future marker development—without attributing causality to specific genes or variants prior to orthogonal and functional validation.

## MATERIAL AND METHODS

### Plant material and genotyping

Four RIL (recombinant inbred line) populations (F7 generation) were considered: Maninjau × Araguaia (M × A), Maninjau × Epagri 108 (M × E), Irat 122 × Araguaia (I × A), and Epagri 108 × Irat 122 (E × I). The process used to obtain these populations is described in Silva et al. (2021; 2022). The design forms a cycle in which each parent participates in two crosses. The raw datasets contained 204 to 293 RILs in the consensus panel and 204 to 293 RILs in the full re-analyses; the number of RILs retained in the mapping core ranged from 127 to 256 for the backbone and from 127 to 247 for the full panel, depending on the quality control criteria specific to each population. The traits evaluated—grain yield, plant height, and days to flowering—were assessed in Goianira and Boa Vista; the experiments are described in Silva et al. (2022).

Genomic DNA was extracted from young leaves using the DNeasy 96 Plant kit (Qiagen, Hilden, Germany). Genome-wide genotyping was performed using DArTseq technology (Diversity Arrays Technology, Canberra, Australia). Markers with a call rate below 0.95 or a minor allele frequency (MAF) below 0.05 were removed. Only markers present in all four populations were retained, resulting in 2,060 SNPs. The full panels contained 15,181 raw SNPs for Maninjau × Araguaia, 5,077 for Maninjau × Epagri 108, 14,365 for Irat 122 × Araguaia, and 9,354 for Epagri 108 × Irat 122. In each population, the SNPs were subjected to filters based on call rate, marker heterozygosity, and minor allele frequency (MAF). The consensus panel workflows predominantly used a MAF of 10% or greater, whereas the full re-analyses adopted a MAF of 5% or greater following the independent freezing of the core set. This discrepancy was controlled, where possible, through paired analyses of the same recombinant inbred lines (RILs) and by projecting the backbone onto the full-panel structure. Physical anchoring of the SNPs was performed by aligning the DArT sequences (TrimmedSequence) to the Nipponbare MSU7/RGAP Release 7 reference genome, accounting for both orientations and correcting the polymorphism position within the aligned sequence. RILs with a call rate below 90% or a residual heterozygosity exceeding the population’s conservative threshold were removed from the core set. As a general rule, a heterozygosity (H) value of 10% or less defined the core; individuals with an extreme number of transitions were identified using a robust criterion based on the median and the Median Absolute Deviation (MAD; Leys et al., 2013). Markers that were locally inconsistent with their flanking markers were removed prior to the final map construction. Parental states were reconstructed based on segregation patterns, historical anchors, and—where available—concordance with the parental genomes.

Co-segregating markers were grouped into bins. “Hard breaks” were maintained in instances where recombination between segments did not support continuity, thereby avoiding artificial interpolation. Distances within blocks were derived from the recombination observed in RILs, with conversion to meiotic recombination using the Haldane mapping function.

### QTL Mapping

QTL analyses were performed separately for each cross, trait, and environment using the R/QTL software (Broman et al., 2003). In the workflows used to reconstruct genotypes along the map via a hidden Markov model (HMM), two parental states were considered. Heterozygous calls and missing data were treated as non-informative observations for the purpose of assigning one of the two parental states. The HMM estimated the probabilities of the parental genotypes across linkage blocks and at intermediate positions within the analysis grid—referred to as “pseudomarkers”—between observed bins. These positions, spaced approximately 1 cM apart, do not correspond to additional genotyped SNPs; rather, they allow for the assessment of QTL support between observed markers. For interval scans, we used Haley–Knott regression (Haley and Knott, 1992). At each map position, the phenotype was regressed on the HMM-estimated probability that a line carried the genotype of one parent. For single-marker scans, the predictor was the observed marker genotype. For scans based on discrete recombination bins, the predictor was the HMM-inferred genotype state (0 or 1). In each single-QTL scan, the LOD score was calculated as (n/2) × log₁₀(RSS₀/RSS₁), where RSS₀ and RSS₁ are the residual sums of squares from the intercept-only model and the model including the genotype predictor, respectively, fitted to the same n lines. Genomic significance thresholds were estimated via permutation, a procedure that determines the empirical distribution of the strongest signal observed during the scan (Churchill and Doerge, 1994). At least 2,000 permutations were performed for each trait.

The stability of QTLs across environments and potential QTL × environment interaction effects were evaluated using components that distinguish the mean performance from the environmental contrast, following the method of Jiang and Zeng (1995). For the lines evaluated in Goianira and Boa Vista, the mean phenotypic values across both environments and the difference (Goianira − Boa Vista) were analyzed, representing the mean component and the environmental contrast, respectively. To assess sensitivity to scale, the values for each trait were standardized separately within each environment using Z-scores—by subtracting the mean and dividing by the standard deviation for the respective environment—after which the analyses of the mean and contrast were repeated. To evaluate the dependency between traits, yield and height were re-analyzed using flowering time as a covariate, and the persistence of the signals following this adjustment was examined. Multilocus models were used to assess the additional statistical contribution of peaks that were nearby or linked, after adjusting for the other loci included in the model, following model-selection approaches for multiple-QTL mapping (Broman and Speed, 2002). Conditional results with nominal significance were treated as complementary analyses and were not interpreted as evidence of genomic significance without comparison against thresholds derived from permutation (Churchill and Doerge, 1994).

### Multi-resolution comparison

The comparison between the backbone and the full-panel datasets was conducted within each cross. A QTL was considered recovered when the peaks coincided, were in very close proximity, or had overlapping support intervals within the same physical region, with a consistent parental effect direction. Comparisons between the backbone and full panels considered the physical positions of QTL peaks, overlap of support intervals, parental effect directions, and significance relative to the permutation-derived genomic threshold for each analysis. These criteria were used to assess whether increased density marker confirmed previously detected regions, improved their localization, or changed the statistical support for individual signals. Where available, matched-RIL analyzes helped distinguish marker-density effects from differences in sample composition. Environmental stability and results from conditional and multilocus analyzes provided complementary evidence for interpreting QTL robustness and independence.

Changes in QTL LOD scores and phenotypic variance explained (PVE) were evaluated considering differences in the number and identity of the RILs included in each analysis. For Maninjau × Araguaia, Maninjau × Epagri 108, and Irat 122 × Araguaia, comparisons controlling for RIL composition were used to assess the contribution of increased marker density. For Epagri 108 × Irat 122, the initial core sets shared 187 RILs, but the comparison focused on whether the major QTLs mapped to corresponding physical regions and showed consistent parental effect directions across panels. Therefore, differences in LOD and PVE in this cross could not be attributed exclusively to marker density. Breakpoints defined by a single RIL were represented as marker-defined recombination-breakpoint intervals rather than statistical confidence intervals. Recombination points were delimited by the informative markers flanking each transition in parental state and were expressed as physical intervals representing the location of the recombination event. These intervals are distinct from QTL support intervals—which are derived from LOD profiles—as they delineate recombination events within the lineages and do not, in themselves, quantify the statistical uncertainty regarding the position of a QTL.

### Integration across backgrounds and prioritization

Recurrence across backgrounds was inferred based on MSU7 position, physical overlap, regional identity, and the direction of the effect from the shared parent. Integration with WGS/BAM data was used to prioritize genes and haplotypes within robust regions. No variant was designated as causal; known genes were treated as functional-positional candidates. Validation via PCR/Sanger sequencing, orthogonal genotyping of recombinants, and functional validation are beyond the scope of this manuscript.

## RESULTS

### Map densification and information gain

The common backbone comprised 2,060 distinct SNPs. Counting each SNP once per cross in which it was mapped, we recorded 3,717 markers across the four backbone maps and 17,323 across the four full-panel maps (a 4.66-fold increase). The total number of recombination bins rose from 2,047 to 4,988 (a 2.44-fold increase), bringing the overall average from 1.82 to 3.47 markers per bin (Table 1). Marker counts increased more than bin counts in every cross, including Maninjau × Araguaia (6.77-fold versus 2.75-fold) and Epagri 108 × Irat 122 (4.21-fold versus 2.94-fold).

**Table 1.**
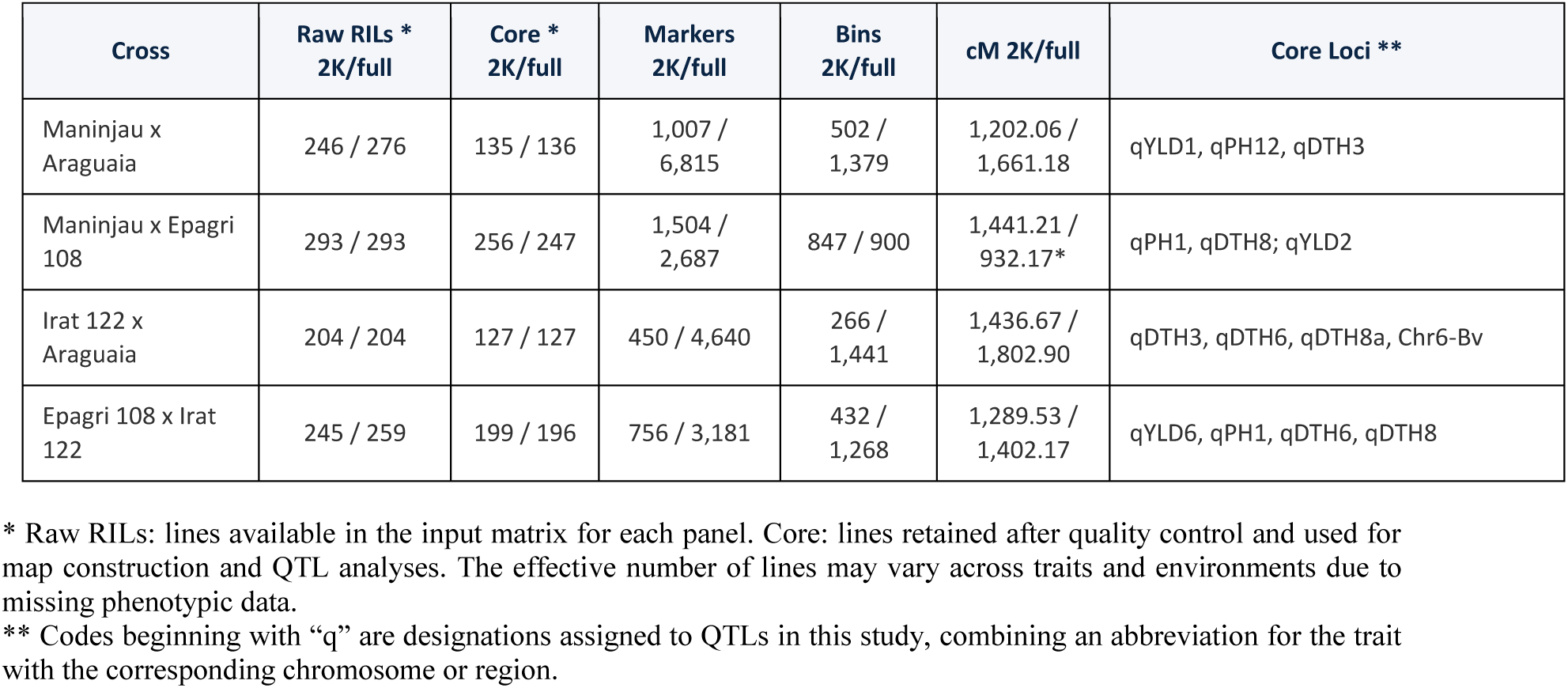
Population sizes, linkage-map characteristics, and selected QTLs in four interconnected rice recombinant inbred line populations analyzed with a common 2,060-SNP backbone and cross-specific full panels. Paired values are reported in backbone/full-panel order. Mapped SNPs are counted separately for each cross, so a backbone SNP can contribute to more than one map.

In Irat 122 × Araguaia, the projected backbone and the full panel detected 2,211 and 4,221 transitions, respectively, in the same 127 RILs, a 90.9% increase. Across the 2,146 paired breakpoints, the median interval width decreased from 1,594.8 to 155.3 kb, a reduction of 90.3%. When the percentage reduction was calculated separately for each pair, the median was 82.1%. Overall, 97.1% of the paired intervals narrowed. In Maninjau × Epagri 108, the number of bins increased from 659 to 889 (34.9%) when the two panels were compared using the same 242 RILs. A separate breakpoint analysis of 247 RILs identified 4,223 comparable events. Their median interval width decreased from 583.0 kb with the backbone to 401.5 kb with the full panel, a reduction of 31.1%. Of these intervals, 1,940 (45.9%) narrowed and 2,283 retained the same width.

### Maninjau x Araguaia: reproduction of the three core QTLs

The three priority QTLs from the backbone were recovered in the full panel. qYLD1 reappeared precisely at Chr1:3,956,269 in the marker/bin scan, with the LOD score increasing from 5.014 to 5.215 and the R² from 15.7% to 18.7%. The Maninjau allele remained associated with higher yield in Goianira, and the signal remained significant in the stable component, even after controlling for flowering time and plant height (Table 2).

**Table 2.** Physical peak positions and LOD scores for grain-yield, plant-height, and flowering-time QTLs detected with the common backbone and cross-specific full panels in Goianira and Boa Vista. Each of the 19 rows compares the two panels for one QTL in one cross and one environment. Peak coordinates give the chromosome and position in base pairs on the Nipponbare MSU7 reference genome.

| **Cross** | **QTL** | **Trait/site** | **2K Peak** | **Full Peak** | **LOD 2K/full** | **Interpretation** |
|---|---|---|---|---|---|---|
| M x A | qYLD1 | Prod_Goi | 1:3,956,269 | 1:3,956,269 | 5.014 / 5.215 | Confirmed; Maninjau favorable |
| M x A | qPH12 | Alt_Goi | 12:22,859,978 | 12:22,430,708 | 6.231 / 5.167 | Confirmed; peak sensitive to the core |
| M x A | qDTH3 | Flo_Goi | 3:1,317,493 | 3:1,317,493 | 9.064 / 11.004 | Strengthened; Araguaia early-maturing |
| M x A | qDTH3 | Flo_Bv | 3:1,279,471 | 3:1,317,493 | 8.481 / 9.125 | Convergence between environments |
| M x E | qPH1 | Alt_Goi | 1:38,426,721 | 1:38,413,000 | 7.646 / 7.215 | Confirmed and stable |
| M x E | qPH1 | Alt_Bv | 1:38,426,721 | 1:38,413,000 | 7.743 / 7.266 | Confirmed and stable |
| M x E | qDTH8 | Flo_Goi | 8:4,322,843 | 8:4,361,000 | 7.177 / 7.856 | Confirmed and stable |
| M x E | qDTH8 | Flo_Bv | 8:4,322,843 | 8:4,361,000 | 5.311 / 5.928 | Confirmed and stable |
| M x E | qYLD2 | Prod_Bv | 2:5,274,193 | 2:5,274,193 | 3.177 / 2.58 | Reclassified; HOLD |
| I x A | qDTH3 | Flo_Goi | 3:1,393,131 | 3:704,454 | 7.179 / 6.25 | Overlapping support levels |
| I x A | qDTH6 | Flo_Goi | 6:9,270,931 | 6:9,112,412 | 4.747 / 4.27 | Recovered; Goianira |
| I x A | qDTH8a | Flo_Goi | 8:4,197,850 | 8:4,138,948 | 8.812 / 9.12 | Recovered; pre-DTH8 |
| I x A | qDTH6-Bv | Flo_Bv | 6:7,535,896 | 6:7,465,338 | 4.242 / 4.17 | Recovered; inverted effect |
| I x A | qPH6-Bv | Alt_Bv | 6:9,908,283 | 6:10,115,609 | 4.512 / 3.91 | Recovered; conditioned |
| E x I | qYLD6 | Prod_Goi | 6:9,094,213 | 6:9,094,213 | 16.51 / 15.90 | Exact reproduction |
| E x I | qPH1 | Alt_Goi | 1:38,391,101 | 1:38,497,531 | 35.45 / 36.93 | Confirmed |
| E x I | qPH1 | Alt_Bv | 1:38,391,101 | 1:38,391,101 | 8.06 / 9.33 | Exact reproduction |
| E x I | qDTH6 | Flo_Goi | 6:9,323,952 | 6:9,270,931 | 12.01 / 11.31 | Refined to *Hd1-CLF* |
| E x I | qDTH8 | Flo_Goi | 8:4,304,925 | 8:4,300,869 | 4.23 / 3.88 | Confirmed |

qDTH3 was the most robust locus in the population. In Goianira, the peak remained at Chr3:1,317,493, with the LOD score increasing from 9.064 to 11.004 and the R² from 26.6% to 35.9%. In Boa Vista, the peak shifted from 1,279,471 to 1,317,493, with an LOD of 9.125 and an R² of 32.2% in the full panel. In both environments, the Araguaia allele accelerated flowering by approximately 6–7 days. Increased marker density refined the recombination-breakpoint interval in RIL 1664 to 90.8 kb, encompassing the *Ef-cd* lncRNA locus and the overlapping protein-coding gene *OsMADS50* (*OsSOC1/DTH3*).

qPH12 remained associated with reduced plant height, with the Araguaia allele conferring the reduction. The peak shifted by approximately 429 kb between the primary backbone and full-panel scans, from Chr12:22,859,978 to Chr12:22,430,708 (Table 2). However, when both marker sets were analyzed using the same 107 RILs, the full panel and the 910 legacy anchors yielded an identical maximum at Chr12:21,918,741. This agreement under matched sample composition suggests that differences in the RILs included in the primary analyzes contributed to the observed peak displacement.

### Maninjau × Epagri 108: confirmation of developmental loci and reclassification of qYLD2

The full-panel qPH1 peak was located in a bin centered at 38.413 Mb on chromosome 1, 13.5 kb from the backbone peak. In the full-panel analysis, the Epagri 108 allele reduced plant height by 14.70 cm in Goianira and 14.46 cm in Boa Vista—values very close to the effects of −14.98 cm and −14.74 cm observed in the backbone analysis. The joint analysis yielded an LOD score of 7.524 and a PVE (proportion of variance explained) of 13.14%, with no significant QTL × environment interaction. The structural span defined by RILs A13SSD0986 and A13SSD0830 was narrowed to 168.5 kb and contains the gene *SD1*/*OsGA20ox2*.

The full-panel qDTH8 peak was located in a bin centered at 4.361 Mb on chromosome 8, 38.4 kb from the backbone peak. qDTH8 was detected in both environments, with the Epagri 108 allele associated with delayed flowering in both cases. The estimated effect was 7.62 days in Goiânia and 5.41 days in Boa Vista; the QTL × environment interaction did not reach genomic significance, although the effect estimates differed between the environments.

In Boa Vista, both panels placed the qYLD2 peak at 5,274,193 bp on chromosome 2, with the Epagri 108 allele associated with higher grain yield. The signal reached genome-wide significance in the backbone scan (LOD 3.18). In the full-panel scan, the LOD score was 2.58, below the genome-wide threshold of 3.05 (permutation P = 0.1274). When both scans were restricted to the same 242 RILs, each yielded a LOD score of 2.594 at that position, below its respective genome-wide threshold. We therefore reclassified qYLD2 as a historical yield signal pending confirmation (Table 2). The full-panel analysis also revealed that the yield signal in Goianira on chromosome 8 was attenuated after adjusting for flowering time.

### Irat 122 x Araguaia: recovered phenological architecture and strong QTL × environment interaction

In Irat 122 × Araguaia, no grain-yield QTL reached genome-wide significance with either marker panel. The full-panel analysis recovered all five priority loci previously identified with the backbone. For each locus, the support intervals from the two panels overlapped. In Goianira, the three flowering-time loci were the recurrent locus qDTH3_rec (qFLO3_Goi in the full panel), qDTH6_Goi (qFLO6_Goi), and qDTH8a_Goi (qFLO8_Goi). In Boa Vista, the full panel recovered the flowering-time locus qDTH6_Bv (qFLO6_Bv) and the plant-height locus qPH6_Bv (qALT6a_Bv).

qFLO3_Goi showed a peak at Chr3:704,454 in the full-panel analysis, with an LOD score of 6.25, a phenotypic variance explained (PVE) of 20.29%, and an effect of −6.61 days (associated with the Araguaia allele). Although there was a 689 kb shift from the previously reported peak at 1,393,131 bp, the support interval (538,353–2,140,481 bp) fully encompassed the original range and included the *Ef-cd* locus.

Increasing marker density narrowed the recombination-breakpoint interval in A15SSD1172 to 53.8 kb, upstream of *Ef-cd*, and identified A15SSD1150 as an additional informative recombinant with a downstream breakpoint. The inferred parental origin at *Ef-cd* differed between the two lines: Araguaia in A15SSD1172 and Irat 122 in A15SSD1150. Together, their recombination patterns bracketed a 247.0-kb candidate region containing *Ef-cd*, providing contrasting materials for targeted genotyping and subsequent genotype–phenotype validation.

On chromosome 6, qFLO6_Goi was located around 9.1 Mb, with the Araguaia allele promoting earlier flowering, whereas qFLO6_Bv was located around 7.47 Mb, with the Araguaia allele delaying flowering. The direct scan of the difference detected qFLO6_GxE at 8.278 Mb, with an LOD of 5.56, PVE of 32.56%, and p_perm = 0.0015. qALT6_GxE was also significant at the 5% level. These results support the existence of components linked to an antagonistic environmental response, rather than a single stable locus.

In Irat 122 × Araguaia, the full-panel scan recovered qDTH8a as qFLO8_Goi, the strongest flowering-time signal in Goianira. The QTL explained 28.15% of the phenotypic variance, and the Araguaia allele was associated with earlier flowering (estimated effect of −7.84 days). The peak mapped to 4,138,948 bp on chromosome 8 (LOD = 9.12), outside the annotated boundaries of DTH8. The full-panel support interval nevertheless encompassed the gene. Backbone markers between 4.30 and 4.34 Mb were monomorphic in this cross, and the available recombinants provided insufficient resolution to distinguish *DTH8* from other regional candidates, including *AP2D9*, *OsGRL17*, and a gene encoding a protein with an Rpb2 domain.

### Epagri 108 × Irat 122: precise replication of qYLD6 and recurrence of qPH1, qDTH6, and qDTH8

In Goianira, both HMM scans placed the qYLD6 peak at the same SNP on chromosome 6 (9,094,213 bp), with the Epagri 108 allele associated with higher grain yield in both panels. The QTL explained 31.75% of the phenotypic variance in the backbone analysis (LOD 16.51) and 32.54% in the full-panel analysis (LOD 15.90).

The qYLD1b signal within the qPH1 region lost statistical support after adjustment for plant height. qPH1 maintained an extreme effect in both Goianira and Boa Vista. In the full-panel analysis, Epagri 108 reduced plant height by 17.29 and 6.78 cm, with LOD scores of 36.93 and 9.33, respectively.

The qDTH6 peak shifted by 53.0 kb on chromosome 6, from 9,323,952 bp in the backbone scan to 9,270,931 bp in the full-panel scan. The Epagri 108 allele was associated with earlier flowering in Goianira in both analyses. In the full panel, SNPs at 9,323,952 and 9,375,785 bp flanked *Hd1* and *OsCLF*. The physical interval around these genes narrowed from 267.8 kb in the backbone to 51.8 kb in the full panel, an 80.6% reduction. The full-panel QTL peak lay outside this interval, 53.0 kb from its lower boundary. The available recombinants did not separate the two genes. The Epagri 108 allele at qDTH8 was associated with later flowering in both panels, and the peak shifted by approximately 4 kb between scans. The two flowering-time loci, qDTH6 on chromosome 6 and qDTH8 on chromosome 8, each remained associated with flowering time after adjustment for the other in the multilocus analysis. In the Epagri 108 × Irat 122 cross, both panels supported qYLD6, qDTH6, and qDTH8, and all three achieved genome-wide significance only in Goianira. qPH1 reached genome-wide significance in both Goianira and Boa Vista.

Across the four crosses, peak positions in the backbone and full-panel scans differed by less than 100 kb in 14 of the 19 comparisons in Table 2 (73.7%), including five exact matches. Each comparison represents one QTL evaluated in one cross and one environment.

### Recurrence between backgrounds

Four QTL regions recurred across crosses sharing a relative. qPH1, encompassing the candidate gene *SD1*, was detected in both Maninjau × Epagri 108 and Epagri 108 × Irat 122, with the Epagri 108 allele consistently reducing plant height in both genetic backgrounds and across both environments. qDTH8/*DTH8* was also centered in Epagri 108. The parent delayed flowering in relation to Maninjau and Irat 122. The effect was stable between Goianira and Boa Vista in the first crossing, but concentrated in Goianira in the second, demonstrating that recurrence of direction does not imply universal environmental stability. qDTH3 was detected in both crosses sharing Araguaia, with the Araguaia allele advancing flowering in each genetic background. In Maninjau × Araguaia, the QTL was detected in both environments and in the analysis of mean flowering time across environments. In Irat 122 × Araguaia, it was detected in Goianira, and the region refined using the full panel included the candidate gene *Ef-cd*. Together, these results support a consistent direction of the Araguaia allelic effect across the two genetic backgrounds, although its detection across environments differed between crosses. qDTH6 was detected in both crosses sharing Irat 122. In Goianira, the Araguaia and Epagri 108 alleles advanced flowering relative to Irat 122 in the 9.1–9.3 Mb region containing the candidate genes *Hd1* and *CLF*. In Irat 122 × Araguaia, a flowering-time QTL was also detected in Boa Vista, but its peak occurred near 7.5 Mb, where the Araguaia allele delayed flowering. The differences in peak position and effect direction suggest a more complex genetic architecture on chromosome 6, potentially involving linked loci with environment-dependent effects, rather than demonstrating an environmental reversal of the same QTL.

Productivity presented a different architecture. qYLD1 occurred in Maninjau x Araguaia, qYLD6 in Epagri 108 x Irat 122 and qYLD2 lost confirmation in the full-panel of Maninjau x Epagri 108; Irat 122 x Araguaia did not present a formal QTL. Therefore, a parental productivity allele with transfer comparable to that observed for height and flowering did not emerge.

### Candidate genes and the limits of inference

Integration of QTL mapping with parental whole-genome sequencing provided complementary evidence for candidate-gene prioritization. QTL positions and recombinant boundaries were compared with gene annotations on the genome reference, while parental sequence variants and, where available, read-level evidence from BAM alignments were examined within the candidate regions. This connected the genomic segments associated with phenotypic variation in the RILs to molecular differences among their parents. The interconnected populations further allowed these candidates to be assessed alongside the direction and recurrence of shared-parent allelic effects across genetic backgrounds. Sequence evidence varied among loci, however, and was used to prioritize hypotheses rather than establish causal variants.

The recurrent qPH1 intervals contained *SD1* (*OsGA20ox2*), and the segment with no observed recombination at qDTH8 contained *DTH8* (*Ghd8*/*Hd5*) (Table 3). At qDTH3, complementary recombinants delimited a 247.0-kb region containing *Ef*-*cd* and the overlapping *OsSOC1* (*OsMADS50*) locus. At qDTH6, parental genome sequences revealed coding variation in *Hd1*. The refined 51.8-kb interval contained both *Hd1* and *OsCLF* (*SDG711*), and the available recombinants did not separate the two genes.

**Table 3.**
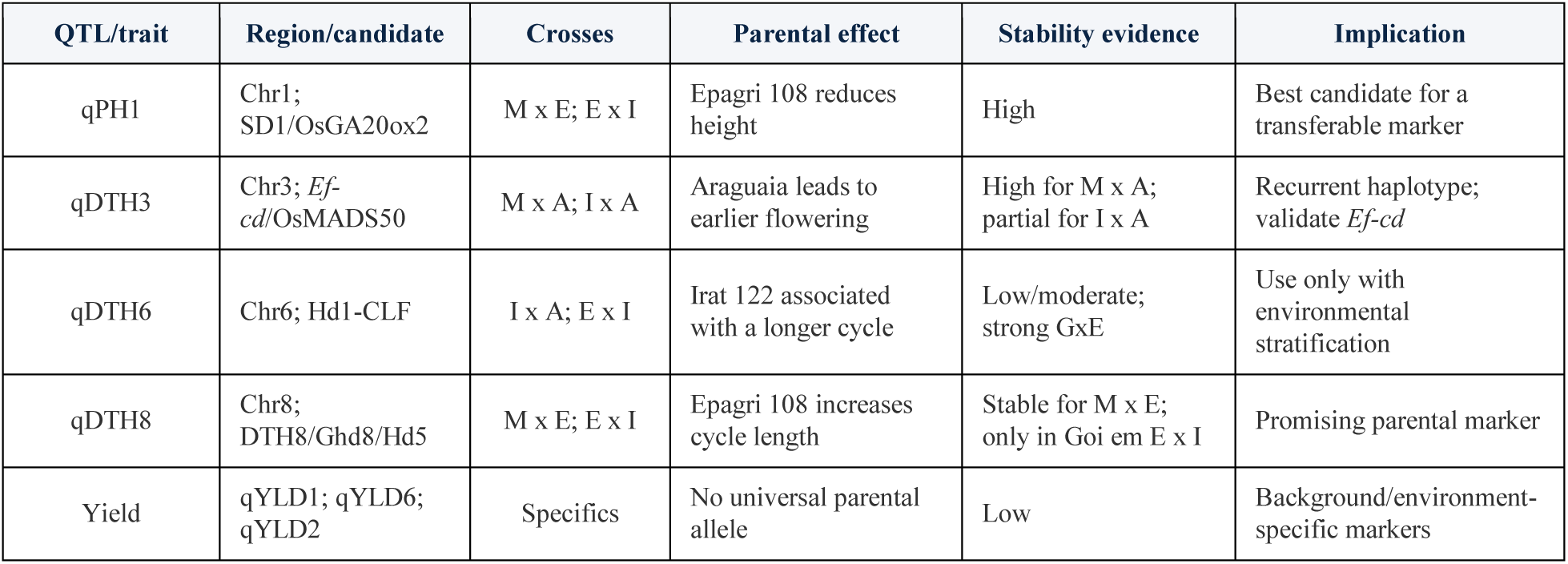
Recurrence of rice QTLs across crosses sharing a parent, consistency of parental effects across environments, and priorities for marker validation. The table summarizes candidate genes and parental effects for four recurrent regions associated with plant height or flowering time, alongside the evidence for grain-yield signals observed in individual crosses.

At qYLD1, reciprocal crossovers in RILs 1405 and 1517 mapped to an 81.7-kb interval on chromosome 1 (3,977,268–4,058,984 bp). *OsBZR5* (qRBG1) lay outside this interval, near the marker-scan peak at 3,956,269 bp. *DNR1* lay within the interval, and the parental origin of its allele remained unresolved in both lines. In qYLD6, the intragenic peak within *OsALDH2b*—along with its independence from flowering time and plant height, and its consistent detection across both analytical resolutions—justifies its prioritization based on both function and position. However, the existence of a documented function or a known parental variant does not prove that the same allele is causal within the populations studied. Consequently, this molecular evidence was used to rank hypotheses and guide future validation efforts, rather than to definitively equate QTLs with causal genes.

## DISCUSSION

### Common backbone and complete panels fulfill different functions

The central methodological contribution of this study is the evaluation of complementary marker-panel strategies in four interconnected rice RIL populations. The common backbone of 2,060 SNPs recovered the major QTL architecture and provided a shared physical framework for comparing QTL regions and parental effect directions across crosses. The cross-specific full panels added information for localizing recombination breakpoints, refining selected regions, and reassessing signals whose support varies between analyses. Their value therefore extended beyond the detection of additional QTLs to improve the interpretation of the regions already identified.

These findings are consistent with previous evidence that increased marker density can improve QTL localization and the resolution of linked loci without necessarily increasing detection power (Stange et al., 2013). Li et al. (2015) further demonstrated that large SNP datasets can be condensed into recombination bins while improving mapping resolution relative to a medium-density map. In the present study, summed across the four linkage maps, marker counts increased 4.66-fold and bin counts increased 2.44-fold. This difference *indica*tes that many additional SNPs described haplotypic segments already represented by other markers. However, the extra markers provided useful local information by narrowing the physical intervals containing inferred recombination breakpoints. Thus, genome-wide redundancy and local gains in resolution can coexist.

The benefit of densification depends on whether additional markers resolve previously indistinguishable genotype patterns or improve the localization of recombination boundaries. Once these boundaries are adequately represented, further markers within an unrecombined segment provide limited additional mapping information. This supports the use of representative markers within bins while retaining denser coverage in regions where it improves resolution. N’Diaye et al. (2017) also showed that retaining excessive numbers of co-segregating markers can contribute to genetic-map expansion under the mapping conditions they examined. The full panels contained more SNPs per bin on average, highlighting the value of grouping markers that co-segregate. A comparison of genetic distances (in cM) between shared markers, using the same RILs and mapping parameters, would help assess whether the denser panels inflated map lengths.

Controlling RIL composition was essential for interpreting differences between marker panels. At qPH12, the primary scans yielded different peak positions, but the full panel and the 910 legacy anchors produced the same maximum when analyzed in the same 107 RILs. This agreement suggests that sample composition contributed to the displacement between the primary scans. In Epagri 108 × Irat 122, the initial core sets shared 187 RILs, but the available comparison did not fully separate marker-density effects from those associated with sample composition. Differences in LOD scores, PVE, and peak positions must therefore be interpreted in light of sample size, RIL identity, genotype frequencies, and analytical settings. Regional overlap, parental effect direction, significance relative to analysis-specific genomic thresholds, and comparisons using the same RILs provide a stronger basis for assessing robustness than an increase in LOD alone.

### Marker densification and complementary analyzes supported a more robust QTL interpretation

Genotyping errors can compromise marker order, agreement between genetic and physical maps, and QTL localization, while missing data can reduce detection power (Wang et al., 2024a; Zhang et al., 2025). Multilocus models and complementary analyzes provide additional means of evaluating whether nearby signals retain statistical support after accounting for other loci (Arends et al., 2010; Bhandari and Lee, 2022). In the present study, binning, HMM-based genotype modeling, conditional analyses, and permutation-based genomic thresholds provided complementary assessments of signal robustness. Their combined use supported a more conservative interpretation of the detected QTLs, although uncertainty remained regarding weaker signals and unresolved linked regions.

Several lost signals support upon reanalysis. In Maninjau × Araguaia, qDTH12 was significant in the single-marker analysis but was not supported by the HMM-based analysis or after conditioning on qDTH3. Likewise, putative QTL × environmental signals on chromosomes 9 and 2 were not recovered in the paired subset. In Maninjau × Epagri 108, qYLD2 retained its peak position and parental effect direction but fell below the genome-wide threshold in the full-panel scan. When both panels were evaluated in the same 242 RILs, they yielded the same peak and LOD score (2.594), below their respective thresholds. The change in significance between the primary scans therefore cannot be attributed to marker density alone. These outcomes justify treating the corresponding signals as provisional. However, loss of significance does not establish the absence of a biological effect: it may also reflect differences in sample composition, statistical power, or model specification. The qPH12 comparison discussed above further illustrates why controlling RIL composition is necessary when interpreting changes between marker panels.

Conversely, several major QTLs showed strong consistency between resolutions. In Epagri 108 × Irat 122, both HMM scans placed qYLD6 at the same SNP in Goianira. The Epagri 108 allele was associated with higher grain yield in both analyses, with LOD scores of 16.51 for the backbone and 15.90 for the full panel. qDTH3 in Maninjau × Araguaia gained support in both environments, while qPH1 and qDTH8 in Maninjau × Epagri 108 retained corresponding positions, parental effect directions, and patterns of detection across environments. In Irat 122 × Araguaia, five previously identified loci were recovered through overlapping support intervals in a full-panel analysis conducted without using the earlier peak positions to guide detection.

Agreement across environments and complementary mapping approaches has also been used to assess QTL reliability in rice (Zheng et al., 2024; Emelin et al., 2024). However, the comparisons presented here reuse the same populations and therefore demonstrate internal consistency across marker resolutions rather than independent validation. Validation with additional genetic materials, as illustrated by the use of reciprocal single-segment substitution lines by Manzoor et al. (2025), remains an important next step. Taken together, the recovery of major loci and the reassessment of weaker signals support a more clearly defined set of QTLs for further investigation. The main gain was greater confidence in the interpretation and prioritization of the mapped regions, rather than an increase in the number of reported loci.

### Developmental QTLs showed greater recurrence across genetic backgrounds than yield QTLs

The Epagri 108 allele at qPH1 reduced plant height in both genetic backgrounds. This recurrent effect is consistent with the role of *SD1* in gibberellin biosynthesis established by Sasaki et al. (2002) and with the effects of *SD1* alleles on plant height reported by Dong et al. (2024). At qDTH3, the Araguaia allele was associated with earlier flowering. Fang et al. (2019) showed that *Ef*-*cd*, a long noncoding RNA transcribed antisense to *OsSOC1*, shortened the time to maturity without reducing yield in the rice lines they evaluated. The candidates at qDTH6 and qDTH8 also have documented roles in photoperiod response. In Goianira, the Epagri 108 allele advanced flowering at qDTH6 and delayed it at qDTH8 in the Epagri 108 × Irat 122 cross. *Hd1* regulates flowering in response to day length (Yano et al., 2000), and a functional *DTH8* allele delayed heading and increased plant height and grain number per panicle under long days (Wei et al., 2010). The effects of these regulators on flowering also depend on the combination of all of them present (Zong et al., 2021). Recurrence and consistent parental effect directions strengthen these regional associations, although causal gene identity and broader transferability remain to be established, particularly for the environmentally dependent qDTH6 region. The opposite effects of the Epagri 108 alleles at qDTH6 and qDTH8— advancing and delaying flowering, respectively—also show that parental contributions must be interpreted locus by locus. Their contributions in multilocus models motivate testing allele combinations for a target flowering window, with the combined effects requiring validation.

Limited recurrence among crosses sharing a relative can have several explanations. A QTL effect is an allelic contrast within a cross: the shared parent may differ functionally from one crossing partner but resemble another at the relevant locus. When functional differences segregate in both crosses, their expression may still depend on genetic interactions and environmental conditions (Wang et al., 2014; Li et al., 2024). Sample size, residual phenotypic variance, recombination patterns, and marker informativeness also influence detection. Consequently, non-detection in a second cross neither establishes biological absence nor demonstrates background dependence. Parental haplotype characterization and targeted validation are needed to distinguish these explanations.

Grain-yield QTLs showed less recurrence. qYLD1 was detected in Maninjau × Araguaia and qYLD6 in Epagri 108 × Irat 122; qYLD2 in Maninjau × Epagri 108 did not retain genome-wide significance in the full-panel analysis, and no yield QTL exceeded the genomic threshold in Irat 122 × Araguaia. This pattern is consistent with the integrated nature of yield, which depends on yield components, architecture, phenology, and growing conditions (Sen et al., 2024), and with reports of environment- and background-dependent yield QTLs in rice (Zhao et al., 2025; Manzoor et al., 2025). Although qYLD6 retained support after adjustment for height and flowering time in Goianira, it was not detected in Boa Vista. Yield-associated markers therefore require validation in the intended genetic background and environment before deployment (Nguyen et al., 2026).

The two positional candidates at qYLD1 suggest different explanations for the observed grain-yield effect. Zhang et al. (2024) showed that a promoter variant of *OsBZR5* (qRBG1) increased grain size and yield in the rice material they studied. Zhang et al. (2021) linked natural variation at *DNR1* to differences in nitrate uptake and assimilation that affected nitrogen-use efficiency and grain yield. In our study, the yield difference between RILs 1405 and 1517 was consistent with their inferred parental haplotypes at *OsBZR5*. The parental origin of *DNR1* remained unresolved in both lines because the gene lay within their crossover interval. Direct genotyping of *DNR1* in these recombinants would show whether either line carries alleles from different parents at the two candidate genes.

The distinct locations of qYLD1 and qYLD6 on chromosomes 1 and 6 make their favorable alleles potential targets for marker-assisted pyramiding. However, detection in separate populations does not establish that their effects will persist or combine advantageously in a common background. Evaluating both all of them individually and together under target growing conditions would determine whether their combination provides an additional yield benefit. Thus, the present results support prioritizing recurrent developmental loci for further transferability testing while treating yield-QTL pyramiding as a hypothesis requiring joint validation.

### Environmental context determines the breeding relevance of a QTL

The chromosome 6 results in Irat 122 × Araguaia illustrate why environmental responses must be interpreted together with physical localization. The Araguaia allele advanced flowering in Goianira at the QTL near 9.1 Mb, whereas it delayed flowering in Boa Vista at a peak near 7.5 Mb. Because these peaks occupy different positions, the opposite effect directions do not establish an environmental reversal of the same causal locus. Instead, they suggest a more complex regional architecture, potentially involving linked loci whose contributions differ between environments. The association of the Irat 122 haplotype with later flowering in the 9.1–9.3 Mb region should therefore be restricted to the crosses and environmental conditions in which that contrast was observed.

In Epagri 108 × Irat 122, qDTH6, qDTH8, and qYLD6 reached genome-wide significance only in Goianira. qPH1 reached this threshold in both Goianira and Boa Vista. By comparison, qPH1 and qDTH8 in Maninjau × Epagri 108 were consistently detected with the same parental effect directions across both environments. These patterns distinguish consistency of detection and effect direction from evidence of similar effect magnitude. Explicit interaction analyzes are needed to determine whether effect differences exceed sampling uncertainty, as demonstrated in multi-location rice RIL experiments by Sakai et al. (2023). Detection in one environment and non-detection in another is insufficient, by itself, to establish QTL × environment interaction; Likewise, an association with mean performance across environments does not establish environmental stability.

For breeding, environmental specificity changes the conditions under which a locus may be useful. Loci with consistent effects across the evaluated environments are promising targets for broader validation, whereas environment-dependent signals may be useful for adaptation to particular growing conditions. Accordingly, qYLD6 remains a candidate for yield improvement in Goianira or comparable environments, subject to validation, while qDTH6 requires evaluation of both the target flowering window and the relevant parental haplotypes. Marker prioritization should therefore consider physical correspondence, effect direction and magnitude, statistical uncertainty, and formal interaction evidence, followed by validation in the intended genetic background and production environment.

Interpreting these environmental responses alongside QTL recurrence, recombinant boundaries, and parental sequence variation provides a stronger basis for prioritizing candidate regions. Consistent parental effect directions across genetic backgrounds support the relevance of recurrent regions, while sequence differences suggest hypotheses about their molecular basis. However, sequence information alone cannot resolve closely linked candidates. At qDTH3, complementary recombinants provide material for investigating the region containing *Ef-cd*, whereas at qDTH6, *Hd1* and *OsCLF*/*SDG711* remain genetically unresolved. These examples show that progress toward causal identification requires both informative molecular variation and genetic material capable of distinguishing competing candidates. The integrated evidence therefore guides targeted validation and marker development, with breeding applications requiring confirmation of favorable haplotypes and their effects in the intended genetic backgrounds and environments.

### Implications for marker-assisted selection

The results support three priorities for marker development and validation. The first comprises qPH1, qDTH3, and qDTH8, which combine recurrence across crosses, consistent parental effect directions, and positional support from established developmental genes. Their candidate regions contain *SD1*/*OsGA20ox2*, involved in gibberellin biosynthesis and height regulation; *Ef-cd*, an antisense lncRNA locus overlapping *OsSOC1*/*OsMADS50*; and *DTH8*/*Ghd8*/*Hd5*, involved in flowering-time regulation. Functional and association studies support the biological relevance of these genes and the importance of their allelic combinations (Fang et al., 2019; Dong et al., 2024; Lee et al., 2025; Zong et al., 2024). These findings justify prioritizing the corresponding regions, although the causal variants underlying the QTLs identified here remain unresolved. A second priority comprises loci supported within particular crosses or environments, including qYLD1, qYLD6, qPH12, and qDTH8a. In Irat 122 × Araguaia, the full-panel support interval for qDTH8a included *DTH8*, leaving its relationship to the recurrent qDTH8 signal unresolved. A third comprises signals requiring further confirmation, such as qYLD2 and the conditional height component qPH6_Bv, which should remain exploratory until additional evidence supports their use.

The qDTH3 results illustrate how improved resolution can guide this process. In Irat 122 × Araguaia, the refined recombination-breakpoint interval of A15SSD1172 and the complementary recombination pattern of A15SSD1150 delimit a 247.0-kb candidate region containing *Ef-cd*. These RILs provide informative material for targeted genotyping and phenotypic evaluation, while the regional boundaries guide the choice of internal and flanking markers. Following confirmation of the parental haplotypes and their associations with flowering time, such markers could support selection of recombinants retaining the desired allele within smaller donor segments. This application is consistent with the staged validation and recombinant-selection strategy described by Platten et al. (2025). The present contribution is a more precise basis for marker development; reduced linkage drag and breeding gains remain outcomes to be tested.

Marker choice should extend beyond the SNP with the highest LOD score. Candidate assays should reliably distinguish the relevant parental haplotypes and reproduce the expected genotypes in parents and informative RILs. Technical validation of genotype calls should then be complemented by testing the marker–trait association in additional segregating or breeding material. For recurrent loci, the two crosses sharing the relevant parent provide an initial framework for evaluating consistency, followed by validation in the intended recipient backgrounds. The desired allele must also be defined by the breeding objective: earlier flowering or reduced height is useful only when it contributes to an appropriate crop duration and plant architecture without unacceptable agronomic penalties.

Environmental targeting is particularly important for qDTH6 and qYLD6. The distinct chromosome 6 flowering signals observed between environments require region-specific haplotype assessment, including further resolution of *Hd1* and *OsCLF*/*SDG711* in the Goianira-associated region. Likewise, qYLD6 should initially be evaluated as a candidate for improving yield under Goianira conditions, with transfer to analogous environments treated as a hypothesis requiring testing. Studies of yield-QTL stability and explicit QTL × environment analysis reinforce the need for this evaluation (Silva et al., 2022; Zhao et al., 2025). Accordingly, validation should establish both the direction and agronomic magnitude of the marker-associated effect under the target production conditions.

Once individual associations have been validated, favorable alleles may be combined to adjust flowering time, plant height, and yield. The separate locations of qYLD1 and qYLD6 make them candidates for testing marker-assisted pyramiding, while the opposite flowering effects of the Epagri 108 alleles at qDTH6 and qDTH8 motivate evaluation of alternative allele combinations. Successful marker-assisted pyramiding in rice demonstrates the practical potential of this approach when marker selection is coupled with phenotypic evaluation (Mohanavel et al., 2024). For the loci identified here, comparing individual and combined haplotypes in a common background would establish whether their effects are retained and whether the combination provides an additional agronomic benefit. This progression from regional prioritization to validated assays, target-environment testing, and evaluation of allele combinations provides a practical route for incorporating the mapping results into rice breeding.

### Recombination-informed panel design and prospective economic efficiency

The results support defining marker density according to the recombination information required for a particular analysis. Although mapped SNP numbers increased 4.66-fold between the backbone and full panels, bin numbers increased only 2.44-fold, *indica*ting substantial redundancy among additional markers. Despite this marker redundancy, the median width of paired breakpoint intervals decreased by 31.1% in Maninjau × Epagri 108 and by 90.3% in Irat 122 × Araguaia. The improvement in recombination-breakpoint localization therefore varied substantially between crosses. This pattern is consistent with improvements in QTL localization at higher marker densities (Stange et al., 2013), the compression of dense SNP datasets into informative recombination bins (Li et al., 2015), and the importance of recombination for mapping resolution (Capilla-Pérez et al., 2024; Wang et al., 2024b). Additional markers can refine the intervals containing inferred recombination breakpoints, but regions lacking informative recombination require additional genetic material to separate linked candidates. Progress toward gene identification therefore depends on combining positional refinement with genetic and functional validation (Manzoor et al., 2025; Sun et al., 2025; Zhang et al., 2026).

Panel performance also depends on the germplasm being analyzed. The C6AIR and 1k-RiCA platforms demonstrated the practical value of intermediate-density genotyping, while showing that marker informativeness varies among genetic groups and parental combinations (Thomson et al., 2017; Arbelaez et al., 2019). For interconnected populations, restricting a panel to polymorphic SNPs in every cross may exclude useful variants specific to individual backgrounds. A more flexible design would combine a common backbone covering the 12 chromosomes with cross-specific modules for poorly represented regions and priority QTLs. Marker selection should consider technical reliability, polymorphism, and coverage of recombination bins across the relevant populations, because bin boundaries are population-dependent. Cross-tailored markers and broader representation of rice genomic diversity provide complementary approaches to these limitations (Steele et al., 2025; Daware et al., 2023).

Parental sequencing can guide variant selection and establish parental allelic states, whereas progeny genotyping is needed to reconstruct inherited segments and locate crossover boundaries. Their complementary roles are illustrated by sequencing-based rice mapping studies (Wang et al., 2024c; Zhang et al., 2026); work in other crops also demonstrates that linkage maps can be constructed with limited parental information when progeny genotypes are available (Cuevas and Vermerris, 2022). For future population series, these findings favor an initial discovery phase with dense parental and progeny information, followed by routine use of a derived backbone and selective densification of informative recombinants or priority intervals. Low-coverage genotyping with imputation offers another potentially economical option, provided that its accuracy is evaluated in the target material (Gorjanc et al., 2017; Sthapit et al., 2025).

This strategy also increases the long-term value of the populations themselves. Producing RILs requires sustained investment in crossing, generation advancement, seed maintenance, phenotyping, and genotyping. Connected populations can support successive analyzes of new traits and environments, as illustrated by multi-cross mapping in rice (Palanog et al., 2023). Here, comparing the backbone with full panels identified information retained by a common marker set and regions benefiting from additional resolution. Reusing the populations and dense reference data can therefore spread the original investment across subsequent biological questions, although the marginal costs of each application still require assessment.

Economic savings remain prospective in the present study because the 2,060-SNP backbone was selected retrospectively from existing data. Routine implementation would require conversion to an appropriate platform: multiplexed targeted sequencing could accommodate the backbone, whereas KASP or TaqMan assays could serve smaller sets of validated priority loci. Reduced AmpSeq panels have demonstrated practical utility in rice breeding (Cerioli et al., 2022), and simulations show that genotyping and phenotyping costs influence resource allocation (Anilkumar and Bernardo, 2026). Any economic comparison should include assay development, validation, sample volume, multiplexing, failed calls, and cross-specific modules. The proposed workflow is consistent with broader efforts to match genotyping platforms to breeding objectives and operational scale (Chu et al., 2026; Bhosale et al., 2025), but its break-even point must be established before a monetary advantage is claimed.

### Implications for crossing design and parental choice

The four populations provided complementary contrasts within and between rice subspecies. Maninjau × Epagri 108 represented an *indica* × *indica* cross, Irat 122 × Araguaia a *japonica* × *japonica* cross, and Maninjau × Araguaia and Epagri 108 × Irat 122 represented *indica* × *japonica* crosses. The two intersubspecific populations yielded the most consistently supported grain-yield QTLs, qYLD1 and qYLD6, respectively. However, the intrasubspecific populations also provided valuable information: Maninjau × Epagri 108 supported consistent height and flowering signals at qPH1 and qDTH8, while Irat 122 × Araguaia revealed multiple flowering-time regions and informative recombinants around qDTH3. Thus, the most useful cross depended on the trait and the allelic contrast being investigated. With only one population representing each intrasubspecific category and two representing the intersubspecific category, differences in parental identity, sample size, recombination, and marker informativeness preclude a general ranking of cross types.

For future crossing blocks, Epagri 108 and Araguaia warrant particular attention as sources of recurrent developmental effects. The Epagri 108 allele at qPH1 reduced height in both Maninjau and Irat 122 backgrounds, supporting its evaluation when reduced stature is a breeding objective. Its flowering-time contribution requires locus-specific selection, because its alleles advanced flowering at qDTH6 but delayed flowering at qDTH8 in the comparisons described here. Araguaia contributed the flowering-advancing qDTH3 allele in both Maninjau and Irat 122 backgrounds and also contributed the height-reducing allele at qPH12 in Maninjau × Araguaia. These findings support investigating Epagri 108 and Araguaia as complementary sources of plant architecture and flowering-time variation, while confirming that the desired haplotypes differ from those already present in recipient lines.

Maninjau and Irat 122 offer additional, more context-dependent opportunities. The favorable Maninjau allele at qYLD1 merits validation in recipient backgrounds selected for yield improvement, whereas the favorable Epagri 108 allele at qYLD6 requires evaluation under Goianira and subsequently other target conditions. Irat 122 provides a contrasting haplotype associated with later flowering in the 9.1–9.3 Mb region in Goianira, making it useful for investigating photoperiod-related variation and for breeding objectives requiring a longer vegetative period, subject to environmental validation. These recommendations concern particular genomic regions and do not establish overall parental superiority or general combining ability. An Epagri 108 × Araguaia cross, which was not evaluated here, is consequently a plausible option for testing the combination of the height-reducing qPH1 haplotype with the flowering-advancing qDTH3 haplotype. Its value would depend on confirming the relevant parental contrasts and evaluating their combined agronomic effects. Alternatively, selected RILs carrying desired haplotypes and suitable agronomic backgrounds could serve as intermediate donors, making further use of the investment in the existing populations. Future crossing decisions should therefore combine subspecies diversity with verified haplotypic complementarity, target-environment performance, and the availability of informative recombinants.

### Conclusions

1. Within the four rice RIL evaluated populations, the consensus backbone of 2,060 SNPs recovered the major QTL architecture and provided a common physical framework for comparing QTL regions and parental effect directions across crosses.
2. Across the four linkage maps, the full panels increased marker counts 4.66-fold and bin counts 2.44-fold. Median breakpoint interval width decreased by 31.1% in Maninjau × Epagri 108 and by 90.3% in Irat 122 × Araguaia, supporting selective marker densification to improve recombination-breakpoint localization.
3. The backbone and full panels recovered the major QTL regions, but some signals were sensitive to the analysis. qYLD2 retained its peak position and parental effect direction but fell below the genome-wide threshold in the full-panel scan. Both panels recovered the same subthreshold peak in the 242 shared RILs. At qPH12, the full panel and the 910 backbone anchors also identified the same peak when compared in the same 107 RILs.
4. The qPH1, qDTH3, qDTH6, and qDTH8 regions recurred across crosses sharing parents, supporting consistent parental allelic contrasts under the conditions in which they were detected. Environmental interpretation required attention to physical position: the opposite flowering effects observed on chromosome 6 in Irat 122 × Araguaia occurred at different peaks and did not establish reversal of the same causal locus.
5. Grain-yield QTLs recurred less frequently across the four populations than did QTLs for plant height and flowering time. The LOD score for qYLD6 exceeded the genome-wide threshold in Goianira and remained below the corresponding threshold in Boa Vista. Markers linked to these grain-yield QTLs therefore require validation in the target genetic background and environment before use in marker-assisted selection.
6. Combining QTL mapping with parental sequencing prioritized candidate genes and informative RILs for validation. Recombinants delimited a 247.0-kb region containing *Ef-cd*, but did not separate *Hd1* from *OsCLF*/*SDG711* at qDTH6. The causal genes and variants remain to be confirmed.
7. Epagri 108 and Araguaia are priority candidates for reducing plant height through qPH1 and advancing flowering through qDTH3, respectively. Their use and the combination of favorable qYLD1 and qYLD6 all require marker validation and confirmation of effects in the target genetic backgrounds and environments.
8. High-density genotyping of founders and initial mapping families can guide the design of a common panel of SNPs representing informative recombination bins. Breeders could use this panel to track validated parental haplotypes in subsequent families and add cross-specific flanking markers at priority loci to select recombinants that retain the desired allele within a shorter donor segment. Because the backbone was derived from existing genotyping data, potential cost savings require prospective evaluation.

## ACKNOWLEDGMENTS

To the Brazilian Agricultural Research Corporation (Embrapa) for financial support (process No. 23.14.01.008.00.00); and to the Goiás State Research Foundation (FAPEG) for the scholarship awarded to the first author (grant No. 202510267001595).

## REFERENCES

Anilkumar, C. & Bernardo, R. (2026). Genomewide Selection in Rice (*Oryza sativa* L.) Breeding: Balancing Cost and Time for Higher Genetic Gain. Plant Breeding, 145(2), 298–305. 10.1111/pbr.70042

Arbelaez, J. D., Dwiyanti, M. S., Tandayu, E., et al. (2019). 1k-RiCA (1K-Rice Custom Amplicon): a novel genotyping amplicon-based SNP assay for genetics and breeding applications in rice. Rice, 12(1), 55. 10.1186/s12284-019-0311-0

Arends, D., Prins, P., Jansen, R. C., et al. (2010). R/qtl: high-throughput multiple QTL mapping. Bioinformatics, 26(23), 2990–2992. 10.1093/bioinformatics/btq565

Bazakos, C., Hanemian, M., Trontin, C., et al. (2017). New Strategies and Tools in Quantitative Genetics: How to Go from the Phenotype to the Genotype. Annual Review of Plant Biology, 68(1), 435–455. 10.1146/annurev-arplant-042916-040820

Bhandari, P. & Lee, T. G. (2022). postQTL: a QTL mapping R workflow to improve the accuracy of true positive loci identification. BMC Research Notes, 15(1), 153. 10.1186/s13104-022-06017-z

Bhosale, S., Thathapalli Prakash, P., Kadaru, S. B., et al. (2025). OneRice breeding framework: An end-to-end system to develop better varieties faster. Crop Science, 65(3), e70087. 10.1002/csc2.70087

Broman, K. W. & Speed, T. P. (2002). A Model Selection Approach for the Identification of Quantitative Trait Loci in Experimental Crosses. Journal of the Royal Statistical Society: Series B (Statistical Methodology), 64(4), 641–656. 10.1111/1467-9868.00354

Broman, K. W., Wu, H., Sen, Ś., et al. (2003). R/qtl: QTL mapping in experimental crosses. Bioinformatics, 19(7), 889–890. 10.1093/bioinformatics/btg112

Capilla-Pérez, L., Solier, V., Gilbault, E., et al. (2024). Enhanced recombination empowers the detection and mapping of Quantitative Trait Loci. Communications Biology, 7(1), 829. 10.1038/s42003-024-06530-w

Cerioli, T., Hernandez, C. O., Angira, B., et al. (2022). Development and validation of an optimized marker set for genomic selection in southern U.S. rice breeding programs. The Plant Genome, 15(3), e20219. 10.1002/tpg2.20219

Chu, H. D., Nguyen, T. Q., Ho, A. Q., et al. (2026). Single-Nucleotide Polymorphism Chip Platforms in Rice (*Oryza sativa*) Genomics and Their Potential for Precision Breeding. International Journal of Plant Biology, 17(9), 81. 10.3390/ijpb17090081

Churchill, G. A. & Doerge, R. W. (1994). Empirical threshold values for quantitative trait mapping. Genetics, 138(3), 963–971. 10.1093/genetics/138.3.963

Collard, B. C. Y. & Mackill, D. J. (2008). Marker-assisted selection: an approach for precision plant breeding in the twenty-first century. Philosophical Transactions of the Royal Society B: Biological Sciences, 363(1491), 557–572. 10.1098/rstb.2007.2170

Cuevas, H. E. & Vermerris, W. (2022). Linkage map construction using limited parental genotypic information. Euphytica, 218, 58. 10.1007/s10681-022-03005-z

Daware, A., Malik, A., Srivastava, R., et al. (2023). Rice Pangenome Genotyping Array: an efficient genotyping solution for pangenome-based accelerated genetic improvement in rice. The Plant Journal, 113(1), 26–46. 10.1111/tpj.16028

Dong, J., Ma, Y., Hu, H., et al. (2024). The Function of *SD1* on Shoot Length and its Pyramiding Effect on Shoot Length and Plant Height in Rice (*Oryza sativa* L.). Rice, 17(1), 21. 10.1186/s12284-024-00699-8

Emelin, M., Qiu, X., Fan, F., et al. (2024). Identification of reliable QTLs and designed QTL breeding for grain shape and milling quality in the reciprocal introgression lines in rice. BMC Plant Biology, 24(1), 38. 10.1186/s12870-023-04707-9

Fang, J., Zhang, F., Wang, H., et al. (2019). *Ef-cd* locus shortens rice maturity duration without yield penalty. Proceedings of the National Academy of Sciences, 116(37), 18717–18722. 10.1073/pnas.1815030116

Fragoso, C. A., Moreno, M., Wang, Z., et al. (2017). Genetic Architecture of a Rice Nested Association Mapping Population. G3: Genes, Genomes, Genetics, 7(6), 1913–1926. 10.1534/g3.117.041608

Gorjanc, G., Dumasy, J.-F., Gonen, S., et al. (2017). Potential of Low-Coverage Genotyping-by-Sequencing and Imputation for Cost-Effective Genomic Selection in Biparental Segregating Populations. Crop Science, 57(3), 1404–1420. 10.2135/cropsci2016.08.0675

Haldane, J. B. S. (1919). The combination of linkage values and the calculation of distances between the loci of linked factors. Journal of Genetics, 8, 299–309.

Haley, C. S. & Knott, S. A. (1992). A simple regression method for mapping quantitative trait loci in line crosses using flanking markers. Heredity, 69(4), 315–324. 10.1038/hdy.1992.131

Jiang, C. & Zeng, Z.-B. (1995). Multiple trait analysis of genetic mapping for quantitative trait loci. Genetics, 140(3), 1111–1127. 10.1093/genetics/140.3.1111

Kosambi, D. D. (1943). The estimation of map distances from recombination values. Annals of Eugenics, 12(1), 172–175. 10.1111/j.1469-1809.1943.tb02321.x

Lee, H., Lee, S. Y., Han, J., et al. (2025). Association of the Allelic Variations in *Hd1*, *OsPRR37*, *Ghd7* and *DTH8* With Major Agronomic Traits in Korean Rice (*Oryza sativa* L.) Cultivars. Plant Breeding, 144(3), 372–386. 10.1111/pbr.13254

Leys, C., Ley, C., Klein, O., et al. (2013). Detecting outliers: Do not use standard deviation around the mean, use absolute deviation around the median. Journal of Experimental Social Psychology, 49(4), 764–766. 10.1016/j.jesp.2013.03.013

Li, C., Li, Y., Bradbury, P. J., et al. (2015). Construction of high-quality recombination maps with low-coverage genomic sequencing for joint linkage analysis in maize. BMC Biology, 13(1), 78. 10.1186/s12915-015-0187-4

Li, W., Boer, M. P., Joosen, R. V. L., et al. (2024). Modeling QTL-by-environment interactions for multi-parent populations. Frontiers in Plant Science, 15, 1410851. 10.3389/fpls.2024.1410851

Malosetti, M., Ribaut, J.-M., & van Eeuwijk, F. A. (2013). The statistical analysis of multi-environment data: modeling genotype-by-environment interaction and its genetic basis. Frontiers in Physiology, 4, 44. 10.3389/fphys.2013.00044

Manzoor, G. A., Yin, C., Zhang, L., et al. (2025). Mapping and Validation of Quantitative Trait Loci on Yield-Related Traits Using Bi-Parental Recombinant Inbred Lines and Reciprocal Single-Segment Substitution Lines in Rice (*Oryza sativa* L.). Plants, 14(1), 43. 10.3390/plants14010043

Mohanavel, V., Muthu, V., Kambale, R., et al. (2024). Marker-assisted breeding accelerates the development of multiple-stress-tolerant rice genotypes adapted to wider environments. Frontiers in Plant Science, 15, 1402368. 10.3389/fpls.2024.1402368

N’Diaye, A., Haile, J. K., Fowler, D. B., et al. (2017). Effect of Co-segregating Markers on High-Density Genetic Maps and Prediction of Map Expansion Using Machine Learning Algorithms. Frontiers in Plant Science, 8, 1434. 10.3389/fpls.2017.01434

Nguyen, T. Q., Ho, A. Q., Chu, H. D., et al. (2026). From Gene-Based Breeding to Haplotype-Based Breeding for Rice (*Oryza sativa*) Improvement in the Genomics Era. International Journal of Molecular Sciences, 27(20), 8972. 10.3390/ijms27208972

Palanog, A. D., Nha, C. T., Descalsota-Empleo, G. I. L., et al. (2023). Molecular dissection of connected rice populations revealed important genomic regions for agronomic and biofortification traits. Frontiers in Plant Science, 14, 1157507. 10.3389/fpls.2023.1157507

Platten, J. D., Henry, A., Chebotarov, D., et al. (2025). The rice Trait Development Pipeline: a systematic framework guiding upstream research for impact in breeding, with examples from root biology. Plant and Soil, 514(1), 29–46. 10.1007/s11104-025-07399-2

Poland, J. A., Brown, P. J., Sorrells, M. E., et al. (2012). Development of High-Density Genetic Maps for Barley and Wheat Using a Novel Two-Enzyme Genotyping-by-Sequencing Approach. PLoS ONE, 7(2), e32253. 10.1371/journal.pone.0032253

Sakai, T., Fujioka, T., Uemura, T., et al. (2023). Whole-genome analysis of recombinant inbred rice lines reveals a quantitative trait locus on chromosome 3 with genotype-by-environment interaction effects. G3: Genes, Genomes, Genetics, 13(6), jkad082. 10.1093/g3journal/jkad082

Sasaki, A., Ashikari, M., Ueguchi-Tanaka, M., et al. (2002). A mutant gibberellin-synthesis gene in rice. Nature, 416(6882), 701–702. 10.1038/416701a

Sen, P., Chatterjee, A., Kumar, D., et al. (2024). Genetic and functional mechanisms of yield-related genes in rice. Acta Physiologiae Plantarum, 46(3), 29. 10.1007/s11738-024-03667-3

Silva, D. R. A., Castro, A. P. de, Cordeiro, A. C. C., et al. (2021). Identificação de variantes alélicas em backgrounds genéticos multiparentais para caracteres associados à produtividade de grão em arroz (Oryza sativa L.). Resumos do 15° Seminário Jovens Talentos, p. 60. Embrapa.

Silva, D. R. A., Mendonça, J. A., Cordeiro, A. C. C., et al. (2022). Identification of stable quantitative trait loci for grain yield in rice. Pesquisa Agropecuária Brasileira, 57, e02812. 10.1590/S1678-3921.pab2022.v57.02812

Spindel, J., Wright, M., Chen, C., et al. (2013). Bridging the genotyping gap: using genotyping by sequencing (GBS) to add high-density SNP markers and new value to traditional bi-parental mapping and breeding populations. Theoretical and Applied Genetics, 126(11), 2699–2716. 10.1007/s00122-013-2166-x

Stange, M., Utz, H. F., Schrag, T. A., et al. (2013). High-density genotyping: an overkill for QTL mapping? Lessons learned from a case study in maize and simulations. Theoretical and Applied Genetics, 126(10), 2563–2574. 10.1007/s00122-013-2155-0

Steele, K., Quinton-Tulloch, M., Vyas, D., et al. (2025). Thousands of trait-specific KASP markers designed for diverse breeding applications in rice (*Oryza sativa*). G3: Genes, Genomes, Genetics, 15(1), jkae251. 10.1093/g3journal/jkae251

Sthapit, S. R., Crain, J., Larson, S., et al. (2025). A low-coverage skim-sequencing and imputation pipeline for genomic selection. The Plant Genome, 18(4), e70139. 10.1002/tpg2.70139

Sun, Z., An, H., Qiu, Z., et al. (2025). Identification of QTLs and a candidate gene affecting rice grain volume via high-density genetic mapping. Frontiers in Plant Science, 16, 1579589. 10.3389/fpls.2025.1579589

Thomson, M. J., Singh, N., Dwiyanti, M. S., et al. (2017). Large-scale deployment of a rice 6 K SNP array for genetics and breeding applications. Rice, 10(1), 40. 10.1186/s12284-017-0181-2

Wang, P., Ma, L., Li, D., et al. (2024b). Fine mapping of the panicle length QTL *qPL5* in rice. Molecular Breeding, 44(2), 6. 10.1007/s11032-024-01443-2

Wang, W., Chen, Z., Yang, Z., et al. (2024c). lcQTH: Rapid quantitative trait mapping by tracing parental haplotypes with ultra-low-coverage sequencing. Plant Communications, 5(10), 101008. 10.1016/j.xplc.2024.101008

Wang, X., Pang, Y., Zhang, J., et al. (2014). Genetic background effects on QTL and QTL × environment interaction for yield and its component traits as revealed by reciprocal introgression lines in rice. The Crop Journal, 2(6), 345–357. 10.1016/j.cj.2014.06.004

Wang, X., Wang, J., Xia, X., et al. (2024a). Effect of genotyping errors on linkage map construction based on repeated chip analysis of two recombinant inbred line populations in wheat (*Triticum aestivum* L.). BMC Plant Biology, 24(1), 306. 10.1186/s12870-024-05005-8

Wei, X., Xu, J., Guo, H., et al. (2010). *DTH8* Suppresses Flowering in Rice, Influencing Plant Height and Yield Potential Simultaneously. Plant Physiology, 153(4), 1747–1758. 10.1104/pp.110.156943

Yano, M., Katayose, Y., Ashikari, M., et al. (2000). *Hd1*, a Major Photoperiod Sensitivity Quantitative Trait Locus in Rice, Is Closely Related to the Arabidopsis Flowering Time Gene *CONSTANS*. The Plant Cell, 12(12), 2473–2483. 10.1105/tpc.12.12.2473

Zaw, H., Raghavan, C., Pocsedio, A., et al. (2019). Exploring genetic architecture of grain yield and quality traits in a 16-way *indica* by *japonica* rice MAGIC global population. Scientific Reports, 9(1), 19605. 10.1038/s41598-019-55357-7

Zhang, H., Wang, J., Cao, S., et al. (2025). Repeated chip analysis for reducing the effect of genotyping errors on gene mapping for two recombinant inbred line populations in wheat (*Triticum aestivum* L.). BMC Plant Biology, 25(1), 1117. 10.1186/s12870-025-07184-4

Zhang, Q., Wu, R., Hong, T., et al. (2024). Natural variation in the promoter of *qRBG1/OsBZR5* underlies enhanced rice yield. Nature Communications, 15(1), 8565. 10.1038/s41467-024-52928-9

Zhang, S., Zhu, L., Shen, C., et al. (2021). Natural allelic variation in a modulator of auxin homeostasis improves grain yield and nitrogen use efficiency in rice. The Plant Cell, 33(3), 566–580. 10.1093/plcell/koaa037

Zhang, Y., Wang, Y., Du, X., et al. (2026). Integrated QTL mapping and candidate gene analysis for yield-related traits and salt tolerance in a rice RIL population. Frontiers in Plant Science, 16, 1711018. 10.3389/fpls.2025.1711018

Zhao, F., Wang, L., & Xu, S. (2025). Identification of QTL-by-environment interaction by controlling polygenic background effect. Journal of Genetics and Genomics, 52(7), 915–926. 10.1016/j.jgg.2025.01.003

Zheng, Y., Ei Khine, E., Mar Thi, K., et al. (2024). Multi-environment BSA-seq using large F₃ populations is able to achieve reliable QTL mapping with high power and resolution: An experimental demonstration in rice. The Crop Journal, 12(2), 549–557. 10.1016/j.cj.2024.01.009

Zong, W., Ren, D., Huang, M., et al. (2021). Strong photoperiod sensitivity is controlled by cooperation and competition among *Hd1*, *Ghd7* and *DTH8* in rice heading. New Phytologist, 229(3), 1635–1649. 10.1111/nph.16946

Zong, W., Song, Y., Xiao, D., et al. (2024). Dominance complementation of parental heading date alleles of *Hd1*, *Ghd7*, *DTH8*, and *PRR37* confers transgressive late maturation in hybrid rice. The Plant Journal, 118(6), 2108–2123. 10.1111/tpj.16732

